# Asynchronous haltere input drives specific wing and head movements in *Drosophila*

**DOI:** 10.1101/2022.09.29.509061

**Authors:** Michael J Rauscher, Jessica L Fox

## Abstract

Halteres are multifunctional mechanosensory organs unique to the true flies (Diptera). A set of reduced hindwings, the halteres beat at the same frequency as the lift-generating forewings and sense inertial forces via mechanosensory campaniform sensilla. Though haltere ablation makes stable flight impossible, the specific role of wing-synchronous input has not been established. Using small iron filings attached to the halteres of tethered flies and an alternating electromagnetic field, we experimentally decoupled the wings and halteres of flying *Drosophila* and observed the resulting changes in wingbeat amplitude and head orientation. We find that asynchronous haltere input results in fast amplitude changes in the wing (“wing hitches”), but does not appreciably move the head. In multi-modal experiments, we find that wing and gaze optomotor responses are disrupted differently by asynchronous input. These effects of wing-asynchronous haltere input suggest that specific sensory information is necessary for maintaining wing amplitude stability and adaptive gaze control.

## Introduction

Animal locomotion, from crawling [1] to swimming [2] to walking [3] to flight [4], typically requires coordinated motion of multiple body segments or appendages. Though this coordination can stem from activity in a central pattern generator, proprioceptive input modulates body motions to adapt each step [5,6]. Specific contraction timing of muscles actuating multiple appendages can significantly change gait, and thus the mechanisms ensuring limb coordination are essential to structuring adaptive locomotion.

In flying flies, wing [7] and haltere [8] proprioceptors provide crucial feedback allowing for active control of each wingstroke to counteract flies’ inherent pitch instability [9]. The halteres are small club-shaped organs which have evolved from hindwings in true flies (Diptera). Long known to be required for stable flight [10], the halteres do not generate significant lift, but instead primarily detect inertial forces using campaniform sensilla at their base [8,11,12]. Afferent neurons from haltere campaniform sensilla entrain their firing to a preferred phase of the haltere stroke cycle [13,14], which in *Drosophila* and most other fly taxa is biomechanically linked to the wing stroke cycle with an antiphase synchrony [15,16].

Stroke-synchronous feedback from halteres is functionally important for wing-steering motoneurons and muscles [8,17,18]. In tethered flight, haltere ablation results in variable wing amplitudes and diminished wing optomotor responses [19–21]. In the blowfly *Calliphora*, some haltere afferents project directly to the motoneuron controlling wing steering muscle wB1 [22]. Stimulation of the haltere nerve can entrain firing of the motoneuron, which is normally more sensitive to wing reafference, if it occurs within a specified time relative to the wing nerve’s firing [23]. Optogenetic activation of haltere muscles resets the firing phase of both the wB1 and wB2 muscles [18]. For these reasons, we hypothesized that asynchronous haltere input is insufficient for stable wing muscle output, and predicted that changing the phase of the haltere’s stroke would cause wing amplitude instabilities distinct from those observed with ablation.

Neck motoneurons also receive input from haltere afferents [24,25], and mediate a similar suite of gaze-stabilizing reflexes [26]. Following bilateral haltere ablation, flies cannot modulate the gain of optomotor gaze responses [27]. However, the head’s movements are driven primarily by vision and not halteres [28]. In some neck motoneurons, spiking responses to visual input are gated, but not modified, by haltere input: they require haltere oscillation but are invariant to haltere frequency [29]. We thus hypothesized that stroke-synchronous haltere information is not necessary for head movements.

Haltere amplitude manipulation results in smooth changes in wingstroke amplitude and gaze direction [28]. However, flies often execute sharp changes in wing stroke amplitude that can take on a variety of time courses. Fast, unilateral “wing hitches” can occur spontaneously [30,31]. These contrast with longer saccades [32,33], which can be either spontaneous or driven by visual input [34–37] and involve coordinated syndirectional motion of both wings and the head [38–41]. Both saccades and wing hitches are distinct outputs of wing and neck motor pathways, and observing how sensory modulation influences them can reveal pathways for processing sensory input [42]

To examine how halteres drive wing and gaze behaviors, we experimentally decoupled wing and haltere movement with both static and dynamic perturbations. By adding a small iron mass to one haltere, we immobilized it, abolishing it as a source of mechanosensory information. Then, using electromagnets, we drove the haltere at specific frequencies, moving it throughout its natural range of motion asynchronously with the wings. An asynchronously-driven haltere increased the likelihood of unilateral wing hitches as well as noise in the wing’s amplitude, but this did not occur in movements of the head. Optomotor responses of both head and wings were significantly diminished. Our results show that asynchronous haltere input is insufficient for wing amplitude stabilization and adaptive optomotor responses, and results in head responses of diminished amplitude.

## Materials and Methods

### Animal care, haltere manipulations, and experimental trials

Animals were reared from a colony of wild-caught *Drosophila melanogaster*. Female flies (3-5 days post-eclosion) were cold-anaesthetized and attached to tungsten pins using UV-curing cement. For magnetic stimulation experiments, a single iron filing was attached to the right haltere with UV-curing cement. Only iron filings of sufficient mass to immobilize the haltere (in excess of ∼12μg, [28]) were used for experiments. For ablation experiments, one or both halteres were completely removed using fine forceps. For filing-treated animals, successive presentations of different haltere stimuli were preceded and followed by control epochs with the magnets disengaged lasting a minimum of 250ms. For each animal, 1-6 epochs from the same experimental condition were drawn for analysis.

### Haltere stimulation

To control haltere kinematics, two iron-core electromagnets were positioned anterior-dorsal and posterior-ventral to the fly (figure 1*a*; [28]). Each electromagnet was powered from a variable power supply (TP1803D, Tekpower, Montclair, CA, USA) switched through a power transistor (D1276A, Panasonic, Kadoma, Japan) and controlled via 5V TTL-level output from a data acquisition board (USB-6434, National Instruments, Austin TX, USA). Pulse trains consisted of 50% duty cycle square wave signals alternating at 25, 36, 50, 72, 100, or 167Hz.

**Figure 1.**
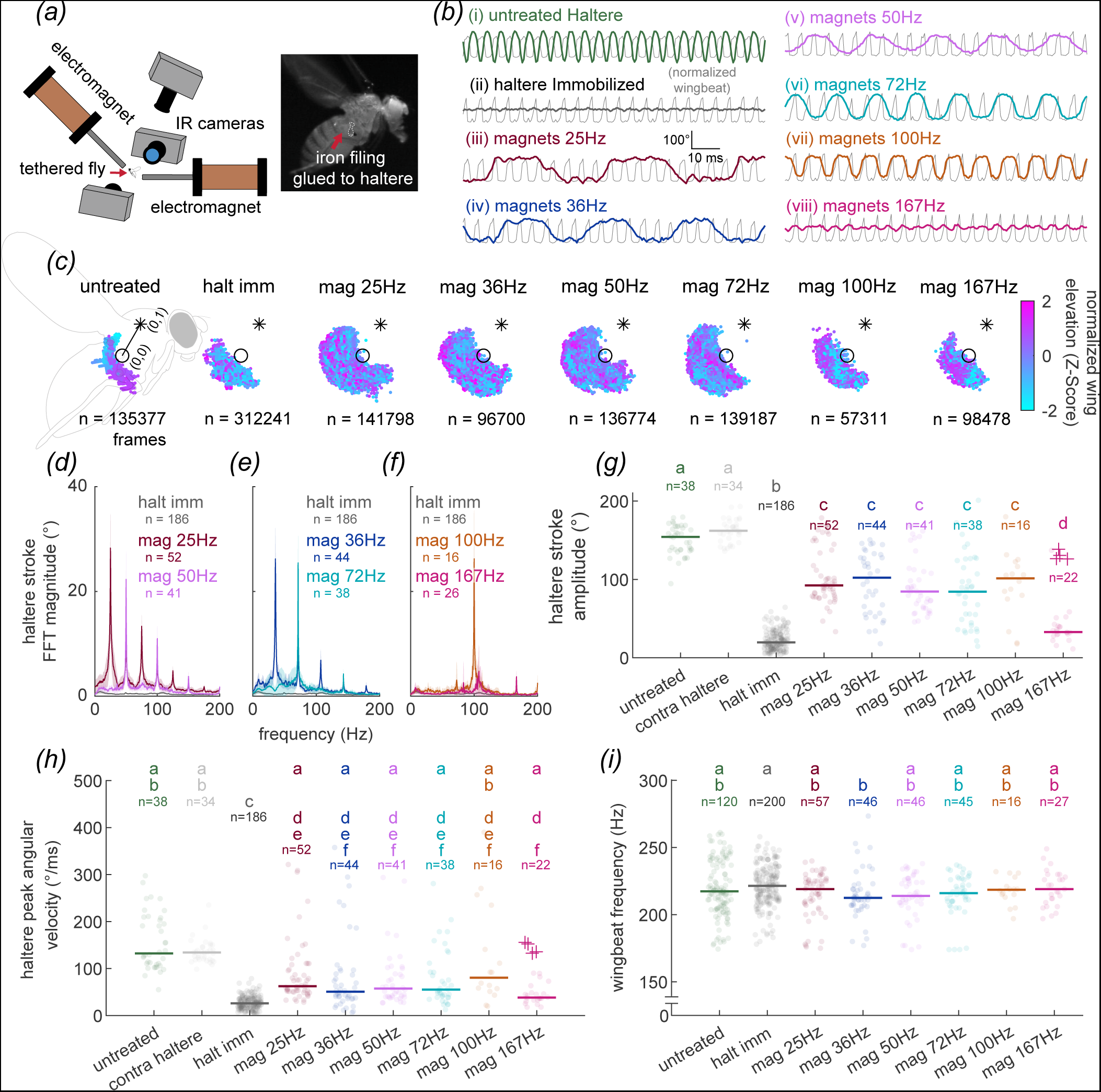
Electromagnetic control of haltere movement asynchronous from the wings. ***(a)*** Schematic of iron filing treatment (top, inset) and electromagnetic stimulation system (bottom). ***(b)*** Representative traces showing movement of untreated haltere at wingbeat frequency (∼200Hz), immobilized haltere treated with iron filing (ii), and actively-driven haltere at a range of frequencies (iii-viii) controlled by electromagnets ***(c)*** Haltere tracking by condition, registered into a normalized body coordinate system defining the haltere base as the origin, represented by circle and the anterior wing base as (0,1), represented by asterisk. Colors reflect the Z-normalized wingstroke elevation at each measurement point. ***(d)*** Mean FFT (95% confidence interval shaded) of all high-speed (1000 or 2000fps) time series with the haltere immobilized (black), driven at 25Hz (red), and 50Hz (pink). ***(e)*** same as *(d)* for 36Hz (blue) and 72Hz (teal) frequencies. ***(f)*** same as *(d)* and *(e)* for 100Hz (orange) and 167Hz (magenta) frequencies. ***(g-i)*** Haltere stroke amplitude *(g)*, peak angular velocity *(h)* and baseline wingbeat frequencies *(i)* observed under each haltere manipulation. Lines show median for each group. 167Hz responses from outlier animal shown with +. Letters denote statistical groupings from multiple Bonferroni-corrected Wilcoxon rank-sum test (5% alpha).

Signals to the magnets were 180° out of phase to produce a dorsal-ventral oscillation at the commanded frequency.

### Visual stimulation

To elicit optomotor responses, we used a wide-field vertical stripe pattern moving in the yaw aspect under the control of a 1Hz triangle wave, presented using a modular LED flight arena [43]. Stripes were one or two pixels wide and pseudo-randomly spaced such that average luminance of the wide-field panorama was 50% [44]. Different pattern velocities were achieved by varying the amplitude of the triangle wave whilst keeping the period constant.

### High-speed video and kinematic measurements

Haltere and wing kinematics were recorded using two synchronized high-speed cameras (TS4 or IL5, Fastec Imaging, San Diego, CA, USA) with sagittal views of the fly from the left and right. Recordings were made at 1000 or 2000fps for all treated animals and the majority of untreated animals; some untreated animals were recorded at 750fps. In each video, the haltere bulb was digitized using the DLTdv8 [45] or DeepLabCut [46] software packages. Wingbeat elevation time series were extracted using a custom MATLAB script. A third camera recording at 100fps (Point Grey Chameleon3, FLIR, Wilsonville, OR, USA) provided a dorsal view to measure head and wing kinematics. Head yaw and wing downstroke envelope kinematic time series were computed using a custom MATLAB program called Flyalyzer (https://github.com/michaelrauscher/flyalyzer; [28]).

### Wing hitch detection and analysis

Wing hitchesCellini et al., 2021 were identified from baseline-subtracted video segments using a custom MATLAB script based around the *findpeaks* function, selecting peaks with a minimum amplitude of 4° (after the saccade detection algorithm in [39]). To capture hitches as described by Götz and Heide [30,31] while excluding saccades, we screened putative hitches for a minimum topographic prominence of 4°, and a maximum duration of 40ms. All detected hitches were used to calculate the hitch rates shown in figure 2*c-d*, figure 4*c-d*, and supplementary figure 2*d-i.* For the event-triggered average (ETA) responses shown in figure 2*e-h* (and supplementary figure 2*a-e*) as well as the cyclic average responses shown in supplementary figure 3, hitches were included only from video segments with concurrent high-speed haltere and wing tracking from the lateral aspect cameras. Each 80ms event-triggered window used to produce the ETA was baseline-subtracted and matched to a control window selected with a randomly drawn frame index from the same video segment. To produce the histograms shown in figure 3*b-g* and the heatmaps shown in supplementary figure 4, data were pooled from all available asynchronously-driven stimulus categories (up to and including 167Hz). Each frame from the 100fps camera was assigned a haltere position and wing instantaneous phase value at frame start by finding the single nearest-match frame recorded on the high-speed lateral aspect cameras (1000 or 2000 fps). To assign a single angular velocity value to each frame from the 100fps camera, the root-mean-square value of the angular velocity time series was computed for all concurrent frames from the lateral aspect camera. Then, the sign (positive or negative) of the lateral aspect frame with the largest absolute value was applied to the result. A graphical summary of this process is included in supplementary figure 4*a*.

**Figure 2.**
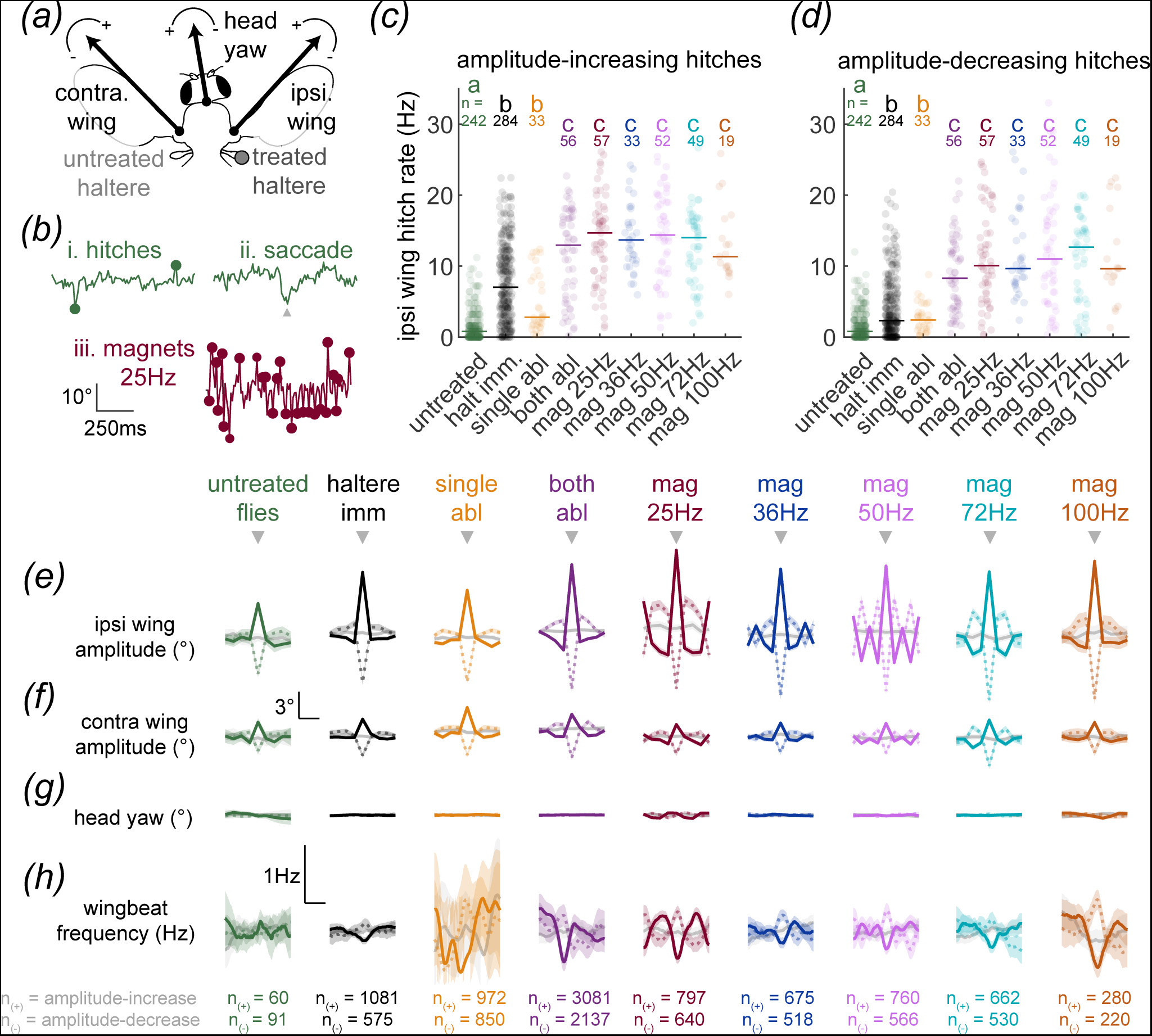
Haltere manipulation increases ipsilateral wing hitch rate. ***(a)*** Schematic of measured kinematic outputs, recorded at 100fps, showing downstroke amplitude (angle of wing envelope leading edge) for each wing and yaw movement of the head. Arcs denote the sign (positive or negative) of clockwise and anticlockwise rotations for each parameter ***(b)*** Example ipsilateral wing amplitude traces showing fast wing hitches from untreated animal (*i*), similar in amplitude but distinct in temporal dynamics from a wing saccade *(ii)*, and increased hitch rate observed in an animal with an asynchronously-driven haltere *(iii)*. Hitches are annotated with filled circles. Saccade with gray triangle ***(c)*** Rate of all ipsilateral wing hitches of increasing amplitude across all haltere treatments (see Methods). Left and right wing responses for symmetrical manipulations (untreated, bilateral ablation) are pooled. Lines show median for each group. Letters denote statistical groupings from multiple Bonferroni-corrected Wilcoxon rank-sum test (5% alpha). ***(d)*** Same as *(c)* but for amplitude-decreasing hitches ***(e)*** Event-triggered average (ETA) responses for each condition, generated from time-aligned, baseline-subtracted kinematic timeseries. Gray triangles show the hitch peak, around which each constitutive video segment of the ETA is indexed, with amplitude-increasing hitch ETAs shown with solid lines and amplitude-decreasing hitch ETAs shown with dashed lines (Shade, 95% confidence interval). Gray traces show controls triggered from random indices within the same time series. ***(f-h)*** Same as *(e)* but for concurrent movements of the contralateral wing *(f)*, head yaw *(g)* and baseline-subtracted instantaneous wingbeat frequency *(h)*.

**Figure 3.**
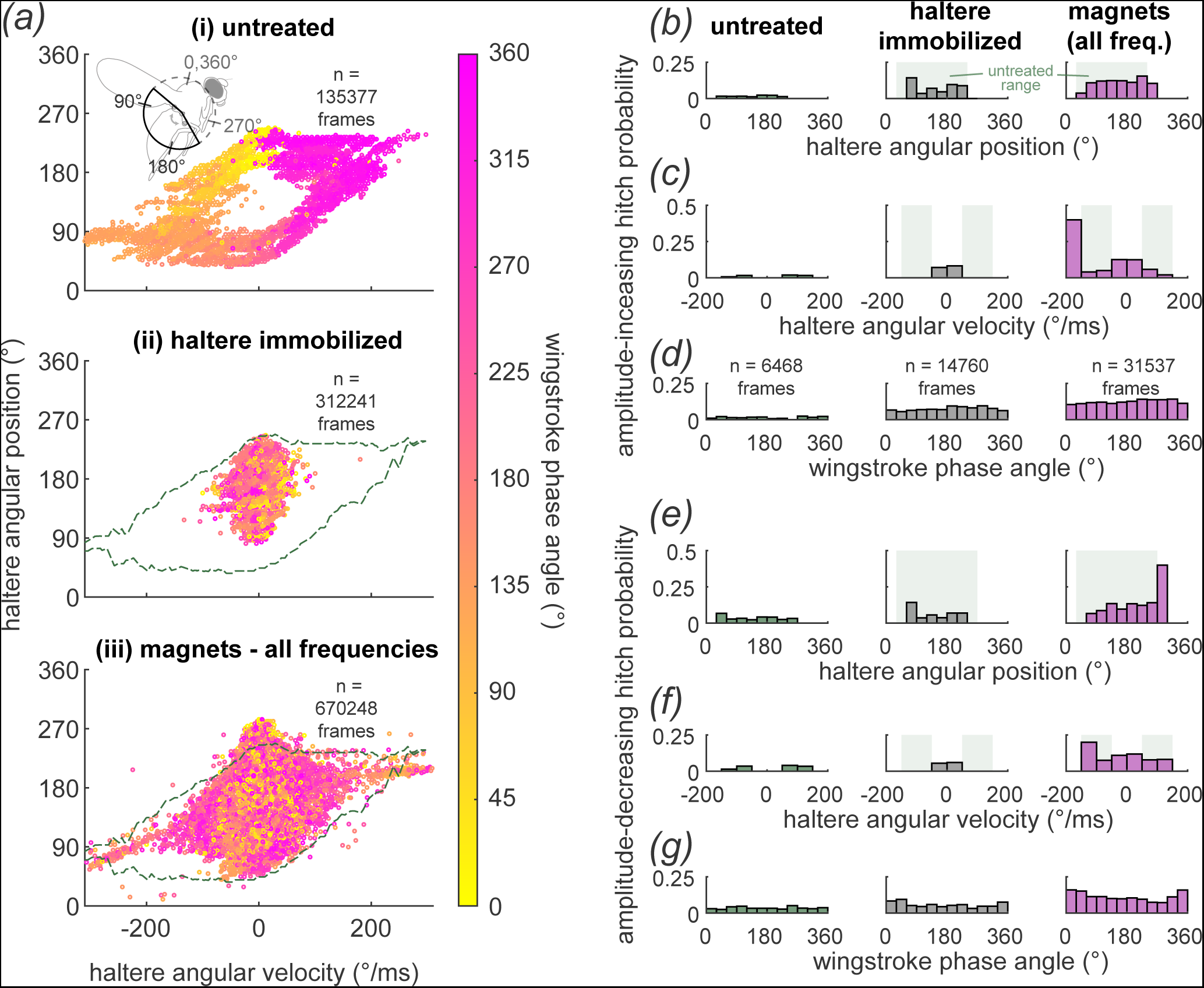
Ipsilateral wing saccades are more likely when the haltere exceeds normal kinematic parameters. ***(a)*** Summary of haltere angular position, angular velocity, and wing instantaneous phase from all high speed data, separated by condition. *(ai)* Untreated animals, showing natural antiphase synchrony of haltere and wings. Inset shows unified body coordinate scheme, with haltere upstrokes typically occupying the 45-90° range and downstrokes typically occupying the 235-270° range *(aii)* Frames from haltere immobilized (magnets OFF) condition. Green dashed line shows extent of the naturalistic parameter space observed in the untreated animal. *(aiii)* High-speed frames aggregated from all magnetic stimulation frequencies, showing broad overlap with the naturalistic range of haltere angular velocities and angular positions. ***(b)*** Histograms of ipsilateral wing amplitude video frames showing probability of amplitude-increasing hitch as a function of haltere angular position at frame onset for each condition. Light green shaded region outlines the natural range of haltere angular positions. ***(c)*** Same as *(b)* but for haltere signed-RMS angular velocity throughout the frame (see Methods). Light green shaded regions outline the naturalistic range of negative (upstroke) and positive (downstroke) angular velocities. ***(d)***. Same as *(b)* but for wing instantaneous phase at frame onset. ***(e-g)*** Same as *(b-d)* but for amplitude-decreasing hitches.

**Figure 4.**
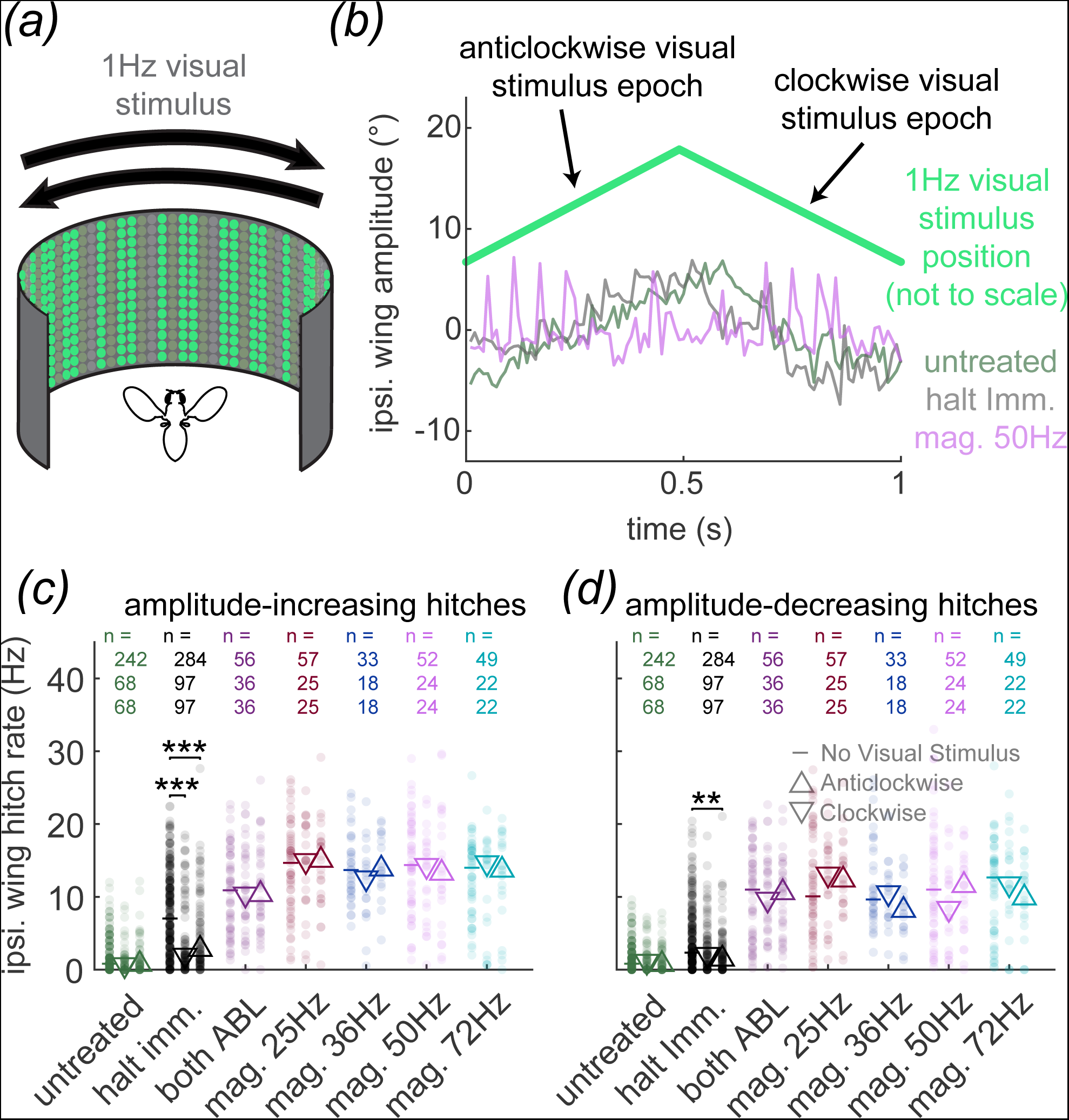
Concurrent visual input does not influence wing hitch rate during most haltere manipulations. ***(a)*** Schematic of visual stimulus, showing irregular stripe pattern moving on an LED panel under the control of a 1Hz triangle-wave with a pattern velocity of 90°/s ***(b)*** Example ipsilateral wing amplitude responses to one cycle of the visual stimulus under several haltere treatment conditions. ***(c)*** Rate of increasing-amplitude ipsilateral wing hitches across visual conditions, showing no change between uniform visual stimulus (reproduced from figure 2), clockwise visual stimulus epochs, and anticlockwise epochs in most haltere treatment conditions. Mean value for each visual stimulus condition represented by dash, upward pointing triangle, and downward pointing triangle respectively. ***(d)*** Same as a *(c)* but for amplitude-decreasing ipsilateral wing hitch rate (** = significance at 1% alpha level, *** = significance at 0.1% alpha level, multiple Bonferroni-corrected Wilcoxon rank-sum test).

### Frequency-domain analysis

For estimation of visual response magnitude (figure 5), kinematic time series were DC-leveled by fitting a least-squares regression line and then subtracting it out to remove linear trends, then zero-padded to 4096 samples. Magnitude estimates from resultant FFTs were then divided by the number of samples in each time series prior to zero-padding to normalize magnitudes for trials of varying length. Only segments longer than 64 samples were used, resulting in a coarsest possible spectral resolution of 1.5Hz.

**Figure 5.**
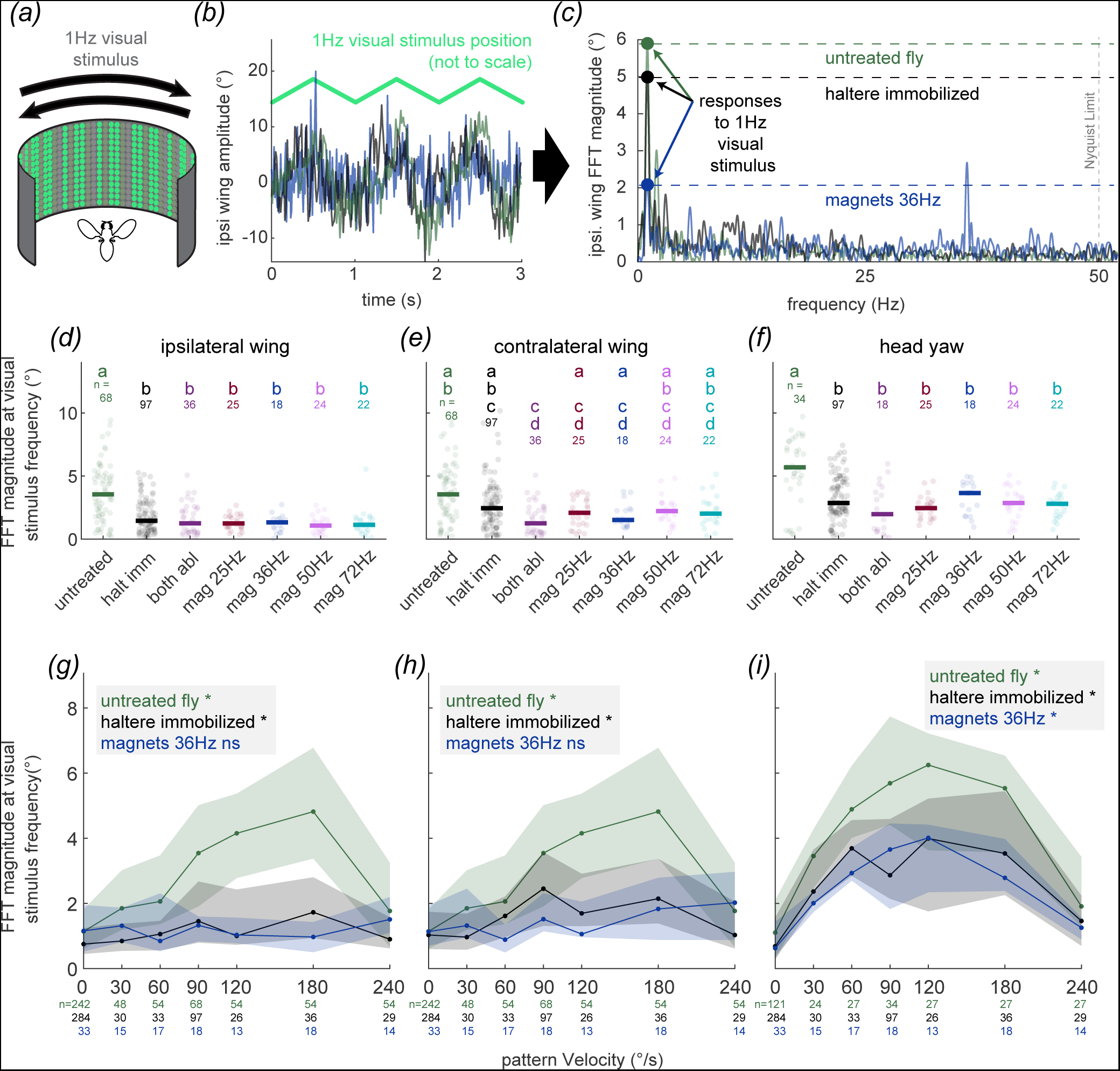
Haltere manipulations influence visually-guided wing and head movements. ***(a)*** Schematic of visual stimulus, as in figure 4A. ***(b)*** Example ipsilateral wing amplitude responses to multiple cycles of the visual stimulus under several haltere treatment conditions. ***(c)*** FFTs of traces from *(b)*, showing peak at the frequency of the visual stimulus. ***(d)*** Ipsilateral wing response amplitude at visual stimulus frequency for all tested haltere treatments. For symmetrical manipulations (untreated, bilateral ablation), left and right wing responses are pooled. Letters denote statistical groupings from multiple Bonferroni-corrected Wilcoxon rank-sum test (5% alpha). ***(e,f)*** Same as *(d)* for contralateral wing and head yaw responses, respectively. Lines show median for each group. ***(g)*** Ipsilateral wing visual response magnitude, as estimated by measuring FFT magnitude at visual stimulus frequency, for untreated flies, haltere-immobilized flies, and 36Hz magnetic stimulation flies. Trendlines connect median data from each pattern velocity, with shaded region showing 1^st^ and 3^rd^ quartiles. Inset, results of Kruskal-Wallis omnibus test indicate whether pattern velocity is a significant predictor of response amplitude within each treatment group at the 5% alpha level (*) or not (ns). ***(h-i)*** Same as *(g)*, but for contralateral wing and head yaw amplitude, respectively.

## Results

### Electromagnetic control of haltere stroke kinematics

To establish magnetic control over one haltere, an iron filing was attached to the bulb (figure 1*a*). With the electromagnets deactivated, this mass immobilized the haltere and prevented the fly from oscillating it. Activation of the electromagnets at 25, 36, 50, 72, 100, and 167 Hz resulted in movement of the haltere at the commanded frequency (figure 1*b-f*, supplemental video 1). This method resulted in haltere movement both within and outside of its naturalistic range (figure 1*c*), with all possible phase relationships to the wing. In all observed trials, the untreated haltere maintained full stroke amplitude (figure 1*g*). At the highest frequency tested (167 Hz), 13 out of 14 flies were unable to maintain a haltere amplitude that was within the range of those observed naturally (figure 1*g*, 22 of 27 replicate trials). We thus excluded 167 Hz trials from further analyses that compared across conditions (see Methods). All conditions achieved peak angular velocities within the range of those observed in untreated animals (figure 1*h*). Wingbeat frequencies were statistically indistinguishable from untreated controls for all treatments (figure 1*i*), showing that manipulations did not impair the gross functioning of the wing motor system.

### Asynchronous haltere movement increases wing hitch rate

Haltere-manipulated flies showed significantly higher rates of wing hitches than untreated control animals, for hitches reflecting wing amplitude increase as well as for hitches of decreasing amplitude (figure 2*a-d*, supplemental video 2). All treatments exhibited higher rates of ipsilateral wing hitches than untreated controls. For both amplitude-increasing hitches (figure 2*c*) as well as amplitude-decreasing hitches (figure 2*d*), dynamic haltere stimulation at any frequency resulted in ipsilateral wing hitch rates comparable to those seen with bilateral haltere ablation, whereas haltere immobilization resulted in intermediate hitch rates comparable to those seen with unilateral ablation.

Were ipsilateral hitches accompanied by coordinated movement of the contralateral wing and head, as seen in saccades? To investigate this, we examined the responses of the contralateral wing and head immediately following an ipsilateral hitch (event-triggered averages; figure 2*e-h*). We observed that under every condition, each ipsilateral wing hitch (figure *2e*, supplemental figure 2*a*) was accompanied by very small changes in the amplitude of the contralateral wing (figure 2*f*, supplemental figure 2*b*) and no meaningful change in head yaw (figure 2*g*, supplemental figure 2*c*). The small changes in contralateral stroke amplitude very rarely met identification criteria for a wing hitch (supplemental figure 2*d-e*). While all treatment groups did exhibit higher rates of contralateral hitches than untreated controls, these hitches were not coordinated with the ipsilateral hitches. Accordingly, the rate of head yaw deviations of equivalent magnitude to a wing hitch was virtually zero under all treatment conditions (supplemental figure 2*e*). Wing and head kinematics during wing hitches therefore did not resemble the coordinated syndirectional movement of the head and wings seen in saccades, emphasizing that these two behaviors are categorically different.[39]

### Haltere kinematic parameters influence hitch initiation

If haltere manipulations directly drive hitches, the rate should increase with increasing stimulus rate. However, wing hitch rate was uniform across stimulation frequencies, and indistinguishable from that seen with bilateral haltere ablation. Despite this, ETAs of the ipsilateral wing amplitude (figure 2*e*) and wing instantaneous frequency (figure 2*h*) oscillated in time with the driving stimulus. As well, fast Fourier transforms (FFTs) of wing and head kinematic time series (supplemental figure 1) showed elevated power at the magnetic stimulation frequency (or its baseband alias, for frequencies which exceeded the Nyquist limit of the kinematics camera). Similarly, stimulus-triggered cyclic averages aligned on each magnetic stimulation pulse showed that hitch probability varied in time with the stimulus for both amplitude-increasing and amplitude-decreasing wing hitches (figure 3*a*). Taken together, these data suggest that hitches occurred at specific times relative to the driving stimulus but were not elicited upon every stroke cycle.

Cyclic averages also illustrated that whereas the treated haltere’s frequency and amplitude were reliably commanded with the electromagnets, the oscillation assumed a diversity of phase relationships with the stimulus across animals and between trials (supplemental figure 3*b-d*). This technical limitation implies that stimulus timing could not necessarily predict specific haltere positions or phases: these had to be measured directly from high-speed video.

After observing phase-locked wing hitches, we examined whether specific haltere kinematic parameters drove hitch initiation. We assessed the haltere’s angular position, angular velocity, and phase relationship to the wingstroke cycle (figure 3*a*). We found that our manipulations resulted in values both within and outside of the range of values observed in untreated animals, allowing us to observe responses to both naturally-occurring stimuli and stimuli that do not occur in flight.

We then created histograms assessing the probability of hitch initiation as a function of each kinematic parameter under each condition, performing separate analyses for amplitude-increasing (figure 3*b-d*) and amplitude-decreasing hitches (figure 3*e-g*). Amplitude-increasing hitch probability was not related to haltere angular position under any condition (figure 3*b*), however amplitude-decreasing hitches were comparatively more likely for the dynamically-stimulated conditions when the haltere was in a downstroke position that exceeded its naturalistic stroke limits (figure 3*e*, right). In untreated animals, the haltere’s angular velocity was almost always high, whether the haltere was moving up or down (figure 3*c*, gray bars); in haltere-immobilized animals, the angular velocity was always low. Hitch probability was also low in these static conditions, although higher in immobilized animals. When we dynamically perturbed the haltere in a velocity range either faster or slower than its natural activity, however, we saw a dramatic increase in amplitude-increasing hitch probability (figure 3*c*, right). For both amplitude-increasing and amplitude-decreasing hitches, there was no relationship between wing phase and hitch probability under any condition (figure 3*d,g*). Multivariate probability heatmaps of multiple kinematic parameters did not reveal any trends over and above those outlined in univariate analysis (supplemental figure 4). Our data therefore show that flies respond to unnatural haltere velocities by initiating ipsilateral wing hitches, with little discernable influence from other measured kinematic parameters.

### Haltere-evoked hitches are not influenced by concurrent visual stimulation

The fly thoracic nervous system integrates inertial sensory information from the halteres with information from the fly’s visual system [25,47,48]. Tethered flies adjust gaze and wingstroke amplitude in response to visual movement [26,49] using a motor pool shared with reflexes driven by the halteres [41,50]. Could visual input influence wing hitches evoked by haltere stimulation?

Concurrently with each haltere treatment, we presented a wide-field pattern of high-contrast vertical stripes with a pattern velocity of 90°/s (figure 4*a-b*, supplemental video 3). We then subdivided each stimulus epoch into periods of clockwise and anticlockwise visual motion (figure 4*a-b*) and calculated hitch rate during each period (as in figure 2). In both visual stimulus directions and for both amplitude-increasing and amplitude-decreasing hitches, visual stimulation did not affect hitch rate under most haltere stimulus conditions (figure 4*c-d*).

Notably, however, concurrent visual stimulation in either direction reduced the rate of amplitude-increasing ipsilateral hitches when the haltere was immobilized (figure 4*c*). For amplitude-decreasing hitches, this was observed only for stimulation in the anticlockwise direction (figure 4*d*). Similar results were observed for the contralateral wing (supplemental figure 5). Our results suggest that descending visual input cannot “rescue” the wing from the effects of asynchronous haltere input.

### Asynchronous haltere movement disrupts wing and head optomotor responses

Halteres adjust the gain of wing and gaze optomotor equilibrium reflexes [20,21,27,41,51]. Is asynchronous input sufficient to drive these? We estimated the optomotor response amplitude (following [51]) by measuring the magnitude of the FFT of each kinematic time series at 1Hz, the frequency of the visual stimulus (figure 5*a-c*). For the ipsilateral wing and head (figure 5*d,f*), all haltere manipulations resulted in smaller visual response amplitudes than those of untreated flies. For the contralateral wing, only the bilateral ablation was statistically distinct (figure 5*e*).

Static haltere manipulations (bilateral ablation and immobilization) impair the ability of flies to adjust their head responses to widefield visual stimuli, locking the fly into a fixed low-amplitude response to all salient pattern velocities [27]. We presented a subset of flies with a series of visual stimuli that moved at a range of pattern velocities, 30-240°/s. Untreated flies exhibited an optomotor tuning curve that peaked at 180°/s for wing responses (figure 5*g-h*, supplemental figure 6*a-b*) and 120°/s for head responses (figure 5*i*, supplemental figure 6*c*), in agreement with previous studies [27,49]. Similar tuning curves were observed for flies with immobilized halteres (figure 5*g-i*) although with significantly lower response amplitudes than untreated flies (supplemental figure 6*d-f*). Under dynamic stimulation (magnets engaged at 36Hz), optomotor tuning was abolished for both contralateral and ipsilateral wings. Visual pattern velocity was not a significant predictor of response amplitude for these flies (figure 5*g-h*). In comparison, head responses showed lower gain relative to untreated controls, at levels comparable to those seen with haltere immobilization (figure 5*i*, supplemental figure 6*c,f*).

## Discussion

For locomotion, coordinated appendages are so important that it is often difficult to force uncoordinated movement: in flying flies, passive thoracic biomechanics restrict halteres and wings to oscillate in antiphase [16]. We used stimuli both within and outside the bounds of natural haltere motions to demonstrate that wing steering reflexes are critically impaired by asynchronous haltere input, with increased ipsilateral wing hitches following “unnatural” haltere movements (figure 2,3). In doing so, we showed that asynchronous haltere input is uniquely deleterious to ipsilateral wing control.

Though we hypothesized that any haltere input would be sufficient for full-amplitude visually guided head movements, we instead observed that asynchronous haltere input results in lower amplitude movements (figure 5). The head’s movements, though driven by a more diverse set of inputs and occurring at lower speeds, still show deleterious effects of asynchronous haltere input. Asynchronous haltere input did not evoke head yaw changes comparable to those seen in the wing (figure 2), congruent with previous findings that abrupt changes in head yaw (as part of a saccade) are primarily evoked by proprioceptive inputs [39]. We show that dynamic haltere perturbations drive “wing hitches”, short-duration changes in amplitude that can occur unilaterally. These hitches may be a small component of the body saccade, a coordinated movement of the head and both wings which is used to reorient the flying fly.

### Asynchronous haltere input increases wing hitches

Asynchronous dynamic haltere input increases the rate of ipsilateral wing hitches (figure 2), particularly when the haltere moves at angular velocities outside of its natural operating range. The halteres are sensitive to the body’s angular velocity [19]; here, we show that they are sensitive to their own angular velocity as well (figure 3). Our experiment with one immobilized haltere shows that hitches are not driven by specific motions of the haltere: loss of one or both halteres also results in an elevated rate of hitches of similar amplitude. Therefore, another possible interpretation is that hitches emerge spontaneously from activity of the wing motor system, and that synchronous haltere input acts to suppress them.

[23][51]Campaniform sensilla are sensitive to compressive and shearing forces [8,52,53], which arise from the haltere’s movement and scale with the bulb’s angular velocity [54]. We might therefore predict that hitches are more or less likely to be initiated as a function of the haltere’s angular position. Our results suggest that amplitude-decreasing hitches are more likely when the haltere is hyperextended past its natural downstroke position, we did not otherwise observe a strong correspondence between haltere angular position and hitch initiation (figure 3).

This suggests, but does not conclusively show, that the halteres suppress spontaneous hitches rather than driving hitches at specific positions. However, the extreme sensitivity of the sensilla [14,55] requires us to consider the possibility that we were not able to stimulate accurately enough to reach a conclusion. The overall effect on flight behavior is the same under both hypotheses: the halteres have a damping effect on wing steering and thus on body rotations, reflecting prior experimental and theoretical work [4,56,57].

### Coordinated movement and haltere influence on the contralateral wing

In previous work [28], manipulating haltere stroke amplitude without compromising haltere-wing synchrony evoked a smooth wing-steering response. Here, responses to haltere perturbation were less stereotyped, but in neither the present nor previous work was there a clear relationship between instantaneous wing phase and haltere-evoked changes in wing kinematic parameters (figure 3). This was surprising given the organization of the wing-steering motor system, a major component of which are tonic muscles which fire once every stroke cycle and control muscle tension by varying firing phase within it [30,58–60]. While our stimulation sampled the entire range of wing/haltere phases, a limitation of our experimental paradigm is that we could not stimulate with a particular phase combination for consecutive stroke cycles, which may be important for the force production properties of these muscles. Another hypothesis is that the untreated contralateral haltere provides enough input to supersede an absent or malfunctioning ipsilateral haltere.

Our findings that asynchronous haltere input increases the rate of contralateral wing hitches independent of movements accompanying ipsilateral hitches (figure 2f, supplemental figure 2b,*d-g*) are consistent with previous findings that each wing receives information from both halteres, but that this information can be used to drive the wings independently. Flies are capable of apparently normal flight following unilateral haltere ablation [61]. In *Calliphora,* input from a single haltere is sufficient to entrain firing of the contralateral first basalare muscle following ablation of the ipsilateral haltere and both wing nerves [17]. Experiments rotating the fly’s entire body showed that both amplitude and frequency components of haltere-wing reflexes are impaired following bilateral—but not unilateral—haltere ablation [19]. Similarly, manipulating the bulb mass of a single haltere produced comparable changes in both clockwise and counterclockwise body saccades [42].

This bilateral integration implicates the contralateral haltere interneurons, or cHINs [24,62], which receive input from haltere afferents and represent the only known anatomical pathway through which haltere information crosses the body axis. Though conserved across fly taxa, these neurons have eluded physiological characterization, leaving questions as to the sensory representations and transformations they mediate *en route* to their motor targets. One possibility is that the predominately unilateral hitches observed in our data might represent “building blocks” that the thoracic nervous system can assemble into more complicated maneuvers such as saccades that coordinate tightly across the body axis.

### Haltere contribution to head and wing optomotor responses

As descending visual inputs concurrently target wing and gaze motoneurons [24,25,63] we also examined multisensory integration of haltere and visual inputs. We show that haltere-wing asynchrony critically impairs the wing optomotor response to a degree comparable with bilateral haltere ablation. While this impairment may be unsurprising for wings, it is less obvious how the head might respond to asynchronous haltere input. Bilateral haltere ablation prevents flies from adjusting the gain of the head optomotor response to different pattern velocities [27]. Here, we show that loss of one haltere (by immobilization) or asynchronous input from one haltere reduces the optomotor response gain, but does not abolish pattern velocity tuning (figure 5).

In some neck motoneurons of quiescent *Calliphora*, an absence of haltere input prevents spiking responses to visual input, but spiking can be restored with haltere movement at any frequency, even as low as 10 Hz [29]. If the halteres are necessary only to provide excitatory drive, then asynchronous input would produce gaze optomotor tuning curves with similar gain to a control. However, we found that asynchronous drive resulted in lower gain.

There are two main hypotheses that could explain why neck motoneurons that can be excited by any haltere input in quiescence fail to produce high-gain visually-guided head movements when stimulated asynchronously in tethered flight. First, wing sensory inputs to neck motoneurons are still unknown, raising the possibility that full neck activity may require both wing and haltere input, like the wing motoneuron mnb1 [22]. However, flies with ablated wings execute high-gain optomotor responses in tethered “flight” ( Mureli et al., 2017).

Second, the “control loop,” in which descending neurons provide excitatory input to haltere muscles, [18,63] may explain this result. Our experiment breaks the control loop by nullifying the relationship between haltere muscle activity and haltere sensory input: visual input to the haltere muscles no longer produces haltere input that can be used to initiate proper reflexes. Both hypotheses could explain our results that visual input cannot “rescue” the wing from high hitch rates (figure 4), that visual tuning is ablated in the ipsilateral wing (figure 5g), and that visual tuning in the head remains intact but at lower gain (figure 5). Future recordings of electrophysiological properties of neck motoneurons under different behavioral conditions and of haltere movement responses to visual inputs could help untangle these possibilities.

## Acknowledgments

We would like to thank Chenxin Bi and Brandon Chun for help with data collection and annotation, as well as Noah DeFino, Jesse Fritz and Lauren Metz for fly care. We would also like to thank Hillel Chiel, Bradley Dickerson, Jeremiah Didion, Alexandra Gurgis, Nicholas Kathman, Kristianna Lea, Mark Willis, Gabriella Wolff, and Alexandra Yarger for helpful commentary.

## Competing interests

No competing interests declared.

## Funding

This work was supported by United States Air Force Office of Scientific Research [FA9550-14-0398 and FA9550-16-1-0165 to JLF]; and National Science Foundation [1754412 to JLF].

## Data Availability

All processed data is available in the associated Dryad repository. Raw video data available upon request. Kinematic time series for some untreated control animals in Figures 1-5 are included with a previously published publicly available dataset [64].

## Notes

### Competing Interest Statement

The authors have declared no competing interest.

### Summary of Updates

Much of the analysis in this manuscript concerns rapid changes in wingbeat amplitude observed during periods of asynchronous haltere stimulation. In prior versions, these were referred to as "saccades." In the present version, these are referred to as "wing hitches" after those observed by Goetz in larger flies (1979 J. Comp. Phys., 1996 JEB). Further discussion of the differences among these two phenomena follow in the text. Analysis in figures 2-4 (as well as in the accompanying supplemental materials) have been substantially revised, separating wing hitches of increasing amplitude from those of decreasing amplitude.

