## Supplemental Materials for "Asynchronous haltere input drives specific wing and head movements in *Drosophila*"

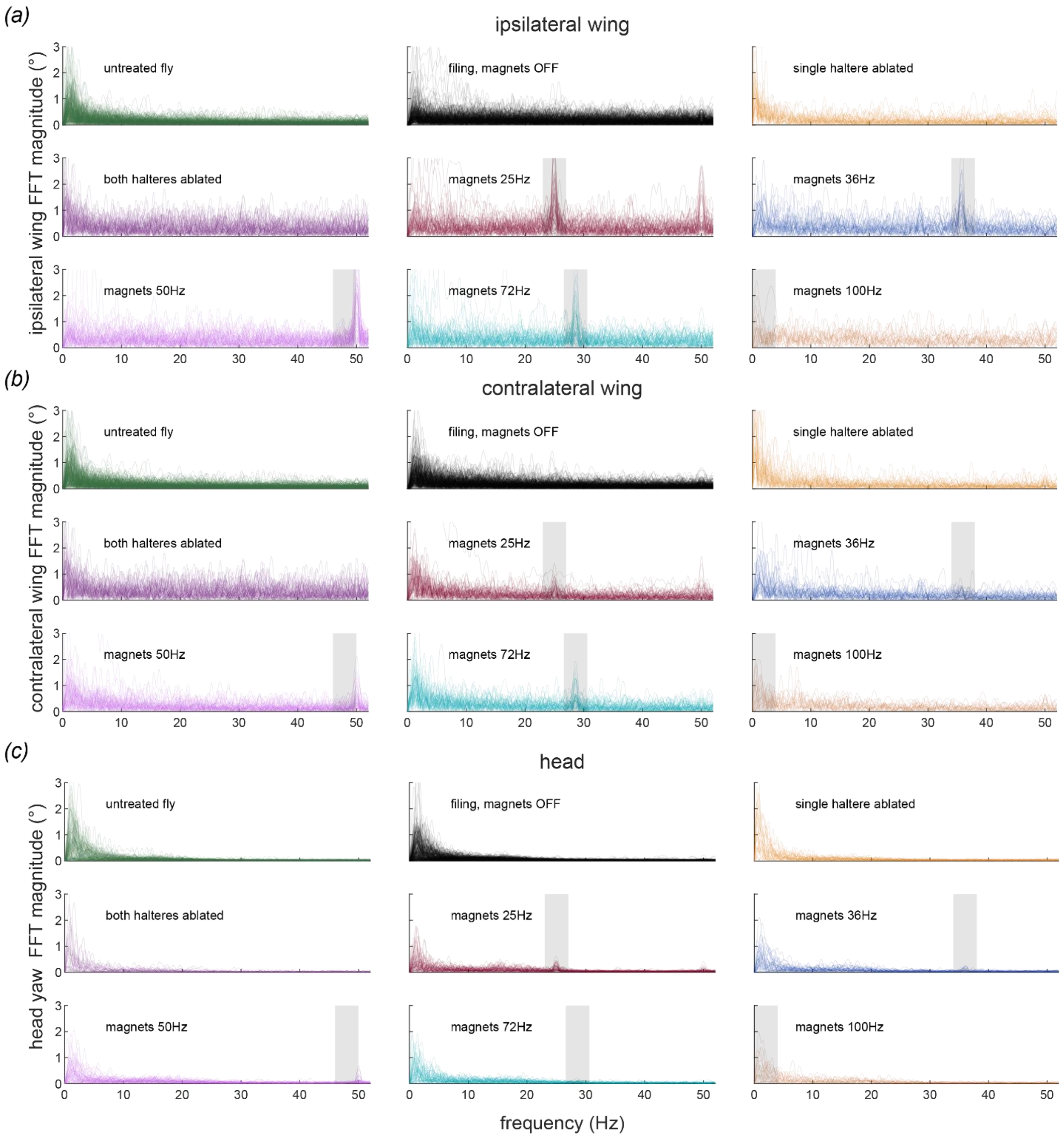


**Supplemental figure 1. Fourier transformations of kinematic outputs downstream of halteres.**
***(a)*** Overlayed FFT for all ipsilateral wing kinematic traces under varying haltere treatment conditions. Shaded regions encompass drive frequency for actively-driven manipulations, or the baseband alias in the case of the 72Hz and 100Hz trials which exceeded the Nyquist limit for the kinematics camera. Left and right wing responses for symmetrical manipulations (untreated, bilateral ablation) are pooled. ***(b,c)*** Same as *(a)*, but for contralateral wing and head responses, respectively.

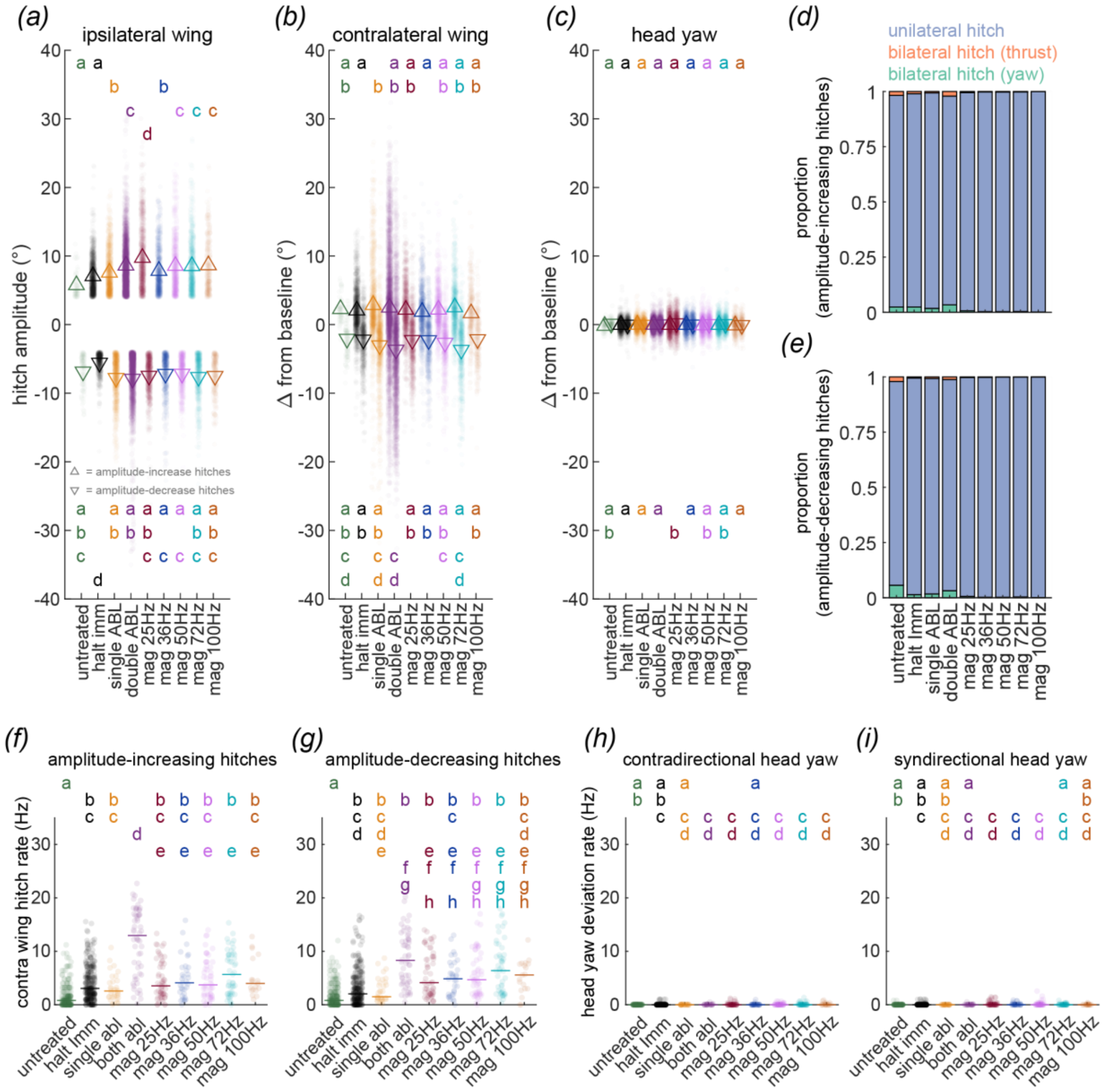


**Supplemental figure 2. Characterization of wing and head kinematic responses to haltere stimulation**
***(a)*** Amplitude of ipsilateral wing hitches as identified in figure 2*f*. Letters denote statistical groupings from one-way ANOVA with post-hoc Tukey test (5% alpha), with upper and lower clusters ascribing to amplitude-increasing and amplitude-decreasing hitches respectively. ***(b)*** Same as *(a)* but showing deviation of concurrent frame of contralateral wingstroke from pre-hitch baseline. ***(c)*** Same as *(b)* but for head yaw, with increasing values showing yaw in the direction away from the treated haltere (contradirectional) and decreasing values showing yaw in the direction of the treated haltere (syndirectional). ***(d)*** Proportion of amplitude-increasing ipsilateral wing hitches that are unilateral (matched time index on contralateral wing meets criteria for hitch detection), bilateral with the contralateral hitch occurring in the same direction ("thrust"), and bilateral with the contralateral hitch occurring in the opposite direction (yaw). ***(e)*** same as *(d)* but for amplitude-decreasing hitches. ***(f)*** Letters denote statistical groupings from multiple Bonferroni-corrected Wilcoxon rank-sum test (5% alpha). ***(h)*** Same as *(f-g)* but for rate of contradirectional head yaw events matching the waveform characteristics of a wing hitch. ***(i)*** Same as *(h)* but for

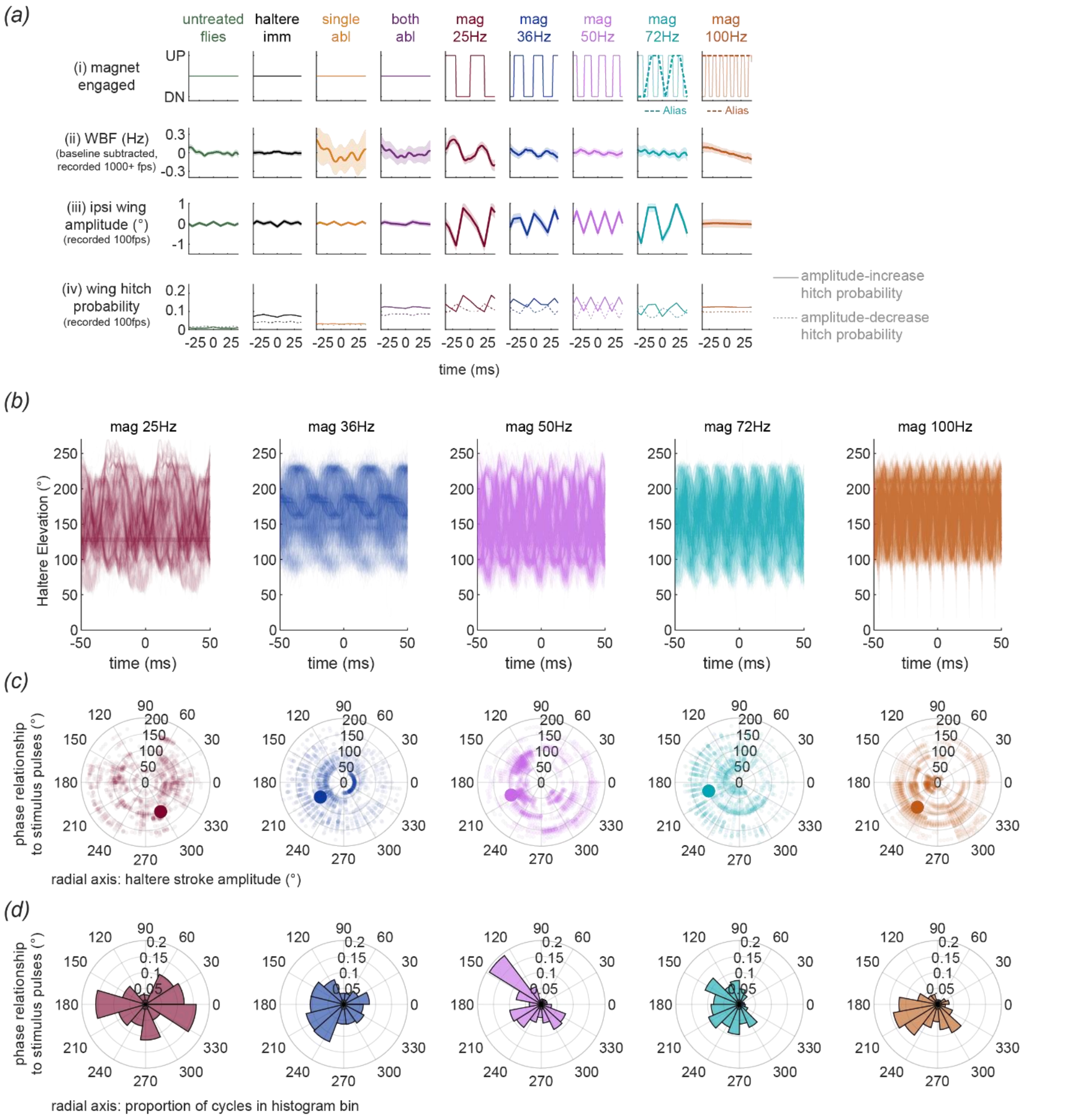


**Supplemental Figure 3. Stimulus-aligned cyclic average responses of dynamic haltere manipulations**
***(a)*** Time-aligned cyclic average responses for selected haltere treatment conditions. *(ai)* Time-course of magnetic control pulses. Dashed line shows baseband alias of the 72Hz and 100Hz magnet pulses on the 100fps overhead camera. *(aii)* Mean baseline-subtracted instantaneous wingbeat frequency (shaded, 95% confidence interval) recorded from 1000-2000fps lateral-aspect camera. ***(aiii)*** Mean baseline-subtracted ipsilateral wing amplitude (shaded, 95% confidence interval). ***(aiv)***. Hitch probability at each time point in *(aiii)*, with amplitude-increasing hitches shown in solid lines and amplitude-decreasing in dashed lines. ***(b)*** Overlayed haltere elevation time series, aligned on each magnetic stimulation pulse cycle. ***(c)*** Polar plots showing amplitude (radial axis) and phase relationship to driving magnetic stimulus (circumferential axis) for each of the cycles depicted in *(b)*. Large marker in each plot shows circular average phase relationship with the driving stimulus and average stroke amplitude. ***(d)*** Polar histogram showing distribution of phase relationships with driving stimulus exhibited by cycles shown in *(b)* and *(c)*. Radial axis shows proportion of cycles in each angular bin.

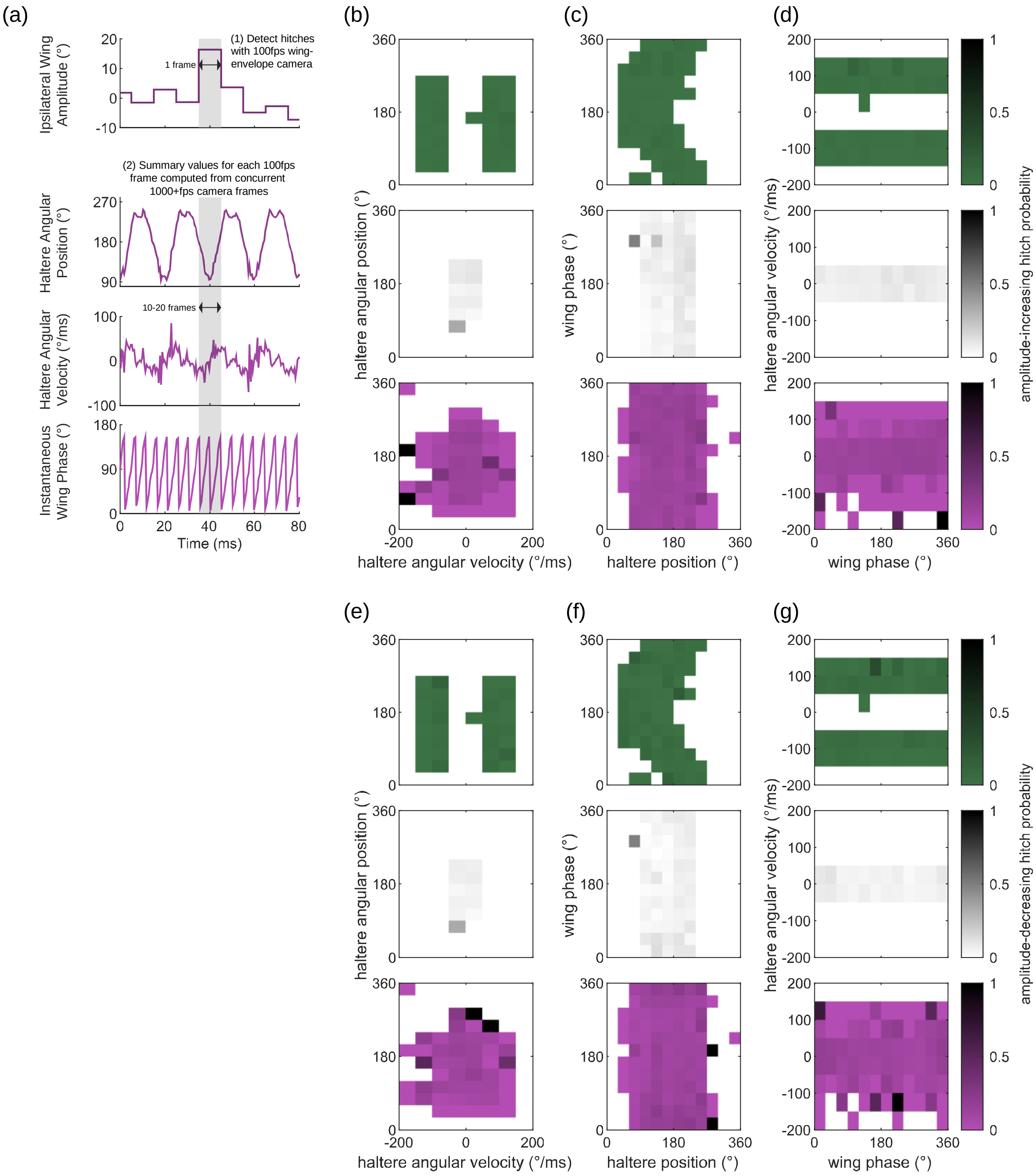


**Supplemental figure 4. Wing hitch probability heatmaps**

***(a)*** Graphical summary of analysis of ipsilateral wing hitches as a function of haltere kinematic parameters, showing wing amplitude trace from 100fps dorsal-view camera with temporally-aligned kinematics from the 1000-2000fps lateral aspect camera (see Methods, *Saccade detection and analysis* for further details). ***(b)*** Heatmap of amplitude-increasing hitch probability as a function of haltere angular position and angular velocity. Binning identical to figure 3b-d. ***(c)*** Same as *(b)* but for haltere angular position and wing instantaneous phase. ***(d)*** Same as *(b)* and *(c)* but for wing instantaneous phase and haltere angular velocity. ***(e-g)*** Same as (b-d) but for hitches of decreasing amplitude.

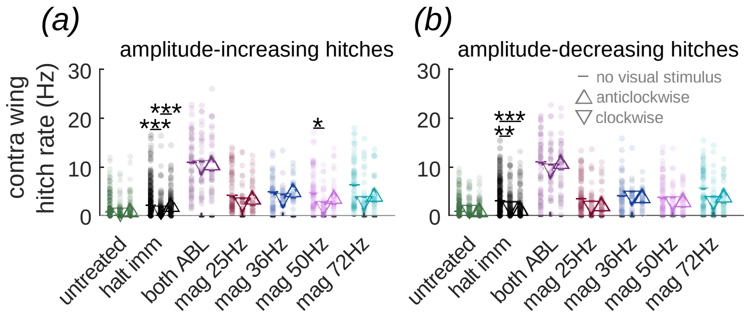


**Supplemental figure 5. Hitch rates of responses to concurrent haltere and visual stimulation**

***(a)*** Rate of increasing-amplitude contralateral wing hitches across visual conditions, showing no change between uniform visual stimulus (reproduced from figure 2), clockwise visual stimulus epochs, and anticlockwise epochs in most haltere treatment conditions. Mean value for each visual stimulus condition represented by dash, upward pointing triangle, and downward pointing triangle respectively. ***(b)*** Same as a *(a)* but for amplitude-decreasing ipsilateral wing hitch rate (pairwise comparisons show outcome of multiple Bonferroni-corrected Wilcoxon rank-sum test. ** = p<1E-2, *** = p<1E-3.)

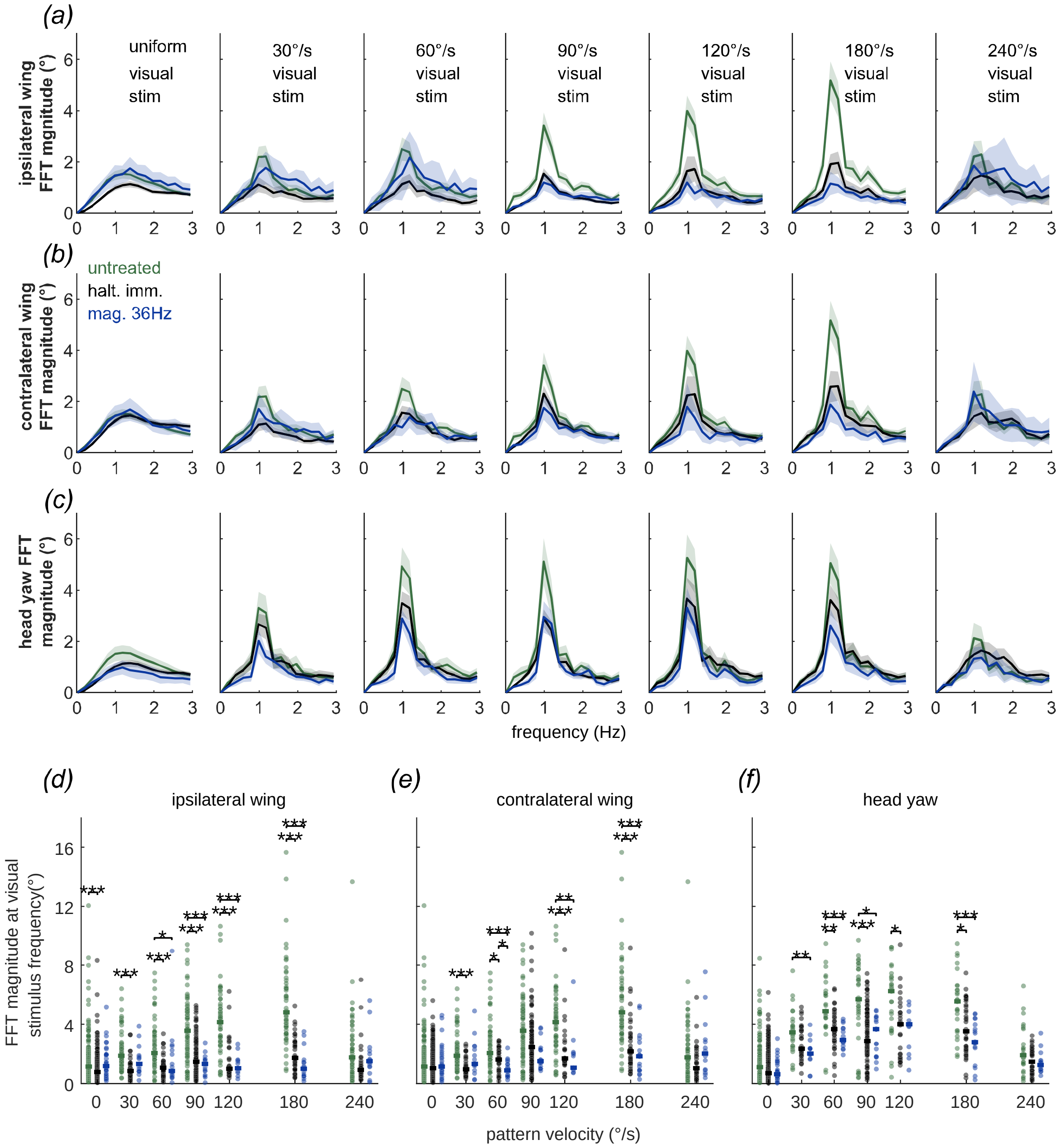


**Supplemental figure 6. Fourier transformations of responses to concurrent haltere and visual stimulation** ***(a)*** Mean FFT and 95% confidence interval (shaded) for ipsilateral wing kinematic traces under varying haltere and visual treatment conditions. ***(b,c)*** Same as *(a)*, but for contralateral wing and head responses respectively. ***(d)*** Magnitude of 1Hz FFT bin for each trial under varying haltere and visual treatment conditions shown in *(a)*, estimating response to visual pattern velocity ***(e,f).*** Same as *(d)*, but for data shown in *(b)* and *(c)* respectively. (pairwise comparisons show outcome of multiple Bonferroni-corrected Wilcoxon rank-sum test *= p<0.05, ** = p<1E-2, *** = p<1E-3. Applicable pairwise comparisons at 90°/s visual pattern velocity are reproduced from figure 5*a*-*c*).

| | Untreated Flies | Contra. Haltere | Halt. Imm. | Mag. 25Hz | Mag. 36Hz | Mag. 50Hz | Mag. 72Hz | Mag. 100Hz | Mag. 167Hz |
|---|---|---|---|---|---|---|---|---|---|
| **Figure 1*c-d*** | | | | | | | | | |
| n = frames | 135377 | | 312241 | 141798 | 96700 | 136774 | 139187 | 57311 | 98478 |
| (flies) | (5) | | (42) | (31) | (27) | (26) | (24) | (18) | (18) |
| **Figure 1*d-f*** | | | | | | | | | |
| n = trials(flies) | | | 186(41) | 52(24) | 44(20) | 41(21) | 38(22) | 16(13) | 26(15) |
| **Figure 1*g-h*** | | | | | | | | | |
| n = trials(flies) | 38(20) | 34(29) | 186(41) | 52(24) | 44(20) | 41(21) | 38(22) | 16(13) | 22(14) |
| **Figure 1*i*** | | | | | | | | | |
| n = trials(flies) | 120(24) | | 200(42) | 57(24) | 46(20) | 46(22) | 45(22) | 16(13) | 27(15) |

| | Untreated Flies | Halt. Imm. | Single Abl. | Both Abl. | Mag. 25Hz | Mag. 36Hz | Mag. 50Hz | Mag. 72Hz | Mag. 100Hz |
|---|---|---|---|---|---|---|---|---|---|
| **Figure 2*c-d*, S1*a,b* *S2f,g*** | | | | | | | | | |
| n = trials(flies) | 242(24) | 284(52) | 33(19) | 56(16) | 57(27) | 33(21) | 52(24) | 49(23) | 19(15) |
| **Figure S1c, S2h,i** | | | | | | | | | |
| n = trials(flies) | 121(24) | 284(52) | 33(19) | 28(16) | 57(27) | 33(21) | 52(24) | 49(23) | 19(15) |
| **Figure 2*e-h*, S2*a-e*** | | | | | | | | | |
| n = amp-inc hitches | 60 | 1081 | 972 | 3081 | 797 | 675 | 760 | 662 | 280 |
| n = amp-dec hitches | 91 | 575 | 850 | 2137 | 640 | 518 | 566 | 530 | 220 |
| (flies) | (4) | (39) | (17) | (15) | (25) | (20) | (22) | (22) | (14) |
| **Fig S3*a-d*** | | | | | | | | | |
| n = cycles | 1600 | 3553 | 7906 | 6328 | 1410 | 1650 | 2695 | 1662 | 2243 |
| (flies) | (5) | (41) | (19) | (15) | (25) | (20) | (22) | (22) | (14) |

| | Untreated Flies | Haltere Immobilized | Magnets (All) |
|---|---|---|---|
| **Fig 3*a*** | | | |
| n = frames (flies) | 135377(5) | 312241(42) | 670248(41) |
| **Figure 3*b-d*, S4*b-g*** | | | |
| n= frames (flies) | 6468(9) | 14760(41) | 31537(38) |

| | Untreated Flies | Both Abl | Halt. Imm. | Mag. 25Hz | Mag. 36Hz | Mag. 50Hz | Mag. 72Hz |
|---|---|---|---|---|---|---|---|
| **Figure 4*c,d, S5a,b*** | | | | | | | |
| No Visual Stim | | | | | | | |
| n = trials(flies) | 242(24) | 56(16) | 284(52) | 57(27) | 33(21) | 52(24) | 49(23) |
| Clockwise Stim | | | | | | | |
| n = trials(flies) | 68(12) | 36(14) | 97(40) | 25(10) | 18(18) | 24(10) | 22(10) |
| Anticlockwise Stim | | | | | | | |
| n = trials(flies) | 68(12) | 36(14) | 97(40) | 25(10) | 18(18) | 24(10) | 22(10) |
| **Figure 5*d,e*** | | | | | | | |
| n = trials(flies) | 68(12) | 36(14) | 97(40) | 25(10) | 18(18) | 24(10) | 22(10) |
| **Figure 5*f*** | | | | | | | |
| n= trials(flies) | 34(12) | 18(14) | 97(40) | 25(10) | 18(18) | 24(10) | 22(10) |

| | No Visual Stimulus | 30°/s V. Stimulus | 60°/s V. Stimulus | 90°/s V. Stimulus | 120°/s V. Stimulus | 180°/s V. Stimulus | 240°/s V. Stimulus |
|---|---|---|---|---|---|---|---|
| **Figure, 5*g-h*, S6*a,b,d,e*** | | | | | | | |
| Untreated Flies | | | | | | | |
| n = trials(flies) | 242(24) | 48(8) | 54(9) | 68(12) | 54(9) | 54(9) | 54(9) |
| Halt. Immobilized | | | | | | | |
| n = trials(flies) | 284(52) | 30(15) | 33(15) | 97(40) | 26(12) | 36(18) | 29(15) |
| Magnets 36Hz | | | | | | | |
| n = trials(flies) | 33(21) | 15(15) | 17(15) | 18(18) | 13(12) | 18(18) | 14(14) |
| **Fig 5*i*,S6*c,f*** | | | | | | | |
| Untreated Flies | | | | | | | |
| n = trials(flies) | 121(24) | 24(8) | 27(9) | 34(12) | 27(9) | 27(9) | 27(9) |
| Halt. Immobilized | | | | | | | |
| n = trials(flies) | 284(52) | 30(15) | 33(15) | 97(40) | 26(12) | 36(18) | 29(15) |
| Magnets 36Hz | | | | | | | |
| n = trials(flies) | 33(21) | 15(15) | 17(15) | 18(18) | 13(12) | 18(18) | 14(14) |

**Supplemental table 1. Number of replicate trials and flies by figure**

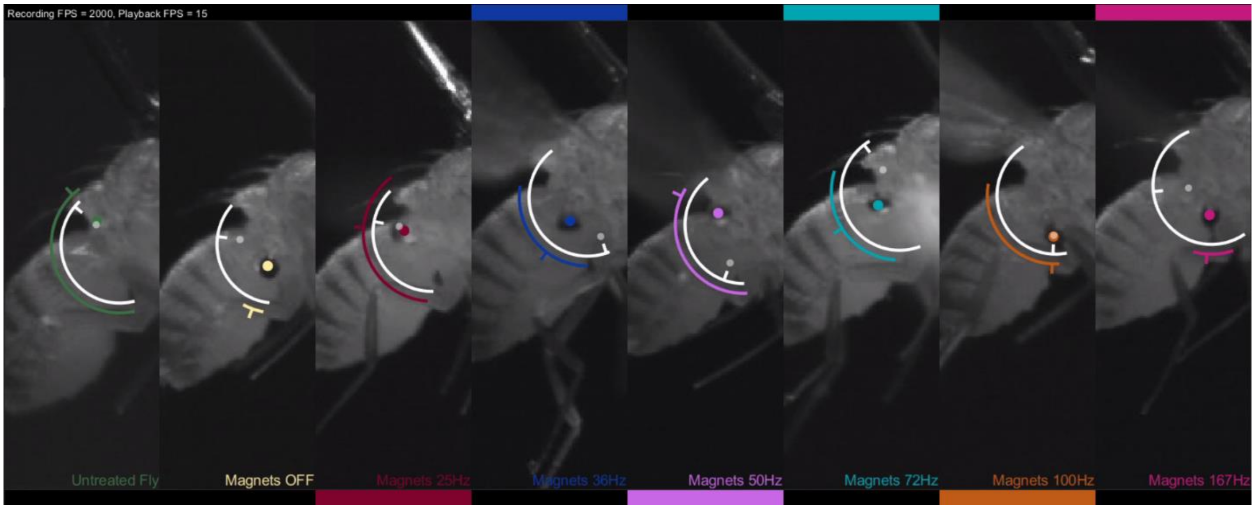


**Supplemental video 1. Electromagnetic control of haltere stroke amplitude**

High speed videos (2000FPS) showing representative magnetic stimulation trials at all tested frequencies. In each video segment, the position of the iron-filing treated haltere bulb is outlined with a colored marker, with an arc of corresponding color and indicator line showing the angular position of the haltere relative to the haltere joint. Position of the untreated opponent haltere (mirrored and reprojected from concurrent high-speed video recording of the contralateral aspect of the fly) is shown with a white marker and angular position with a matching white arc and indicator. Colored boxes above and below each applicable video panel show control pulses energizing the upper and lower electromagnets respectively.

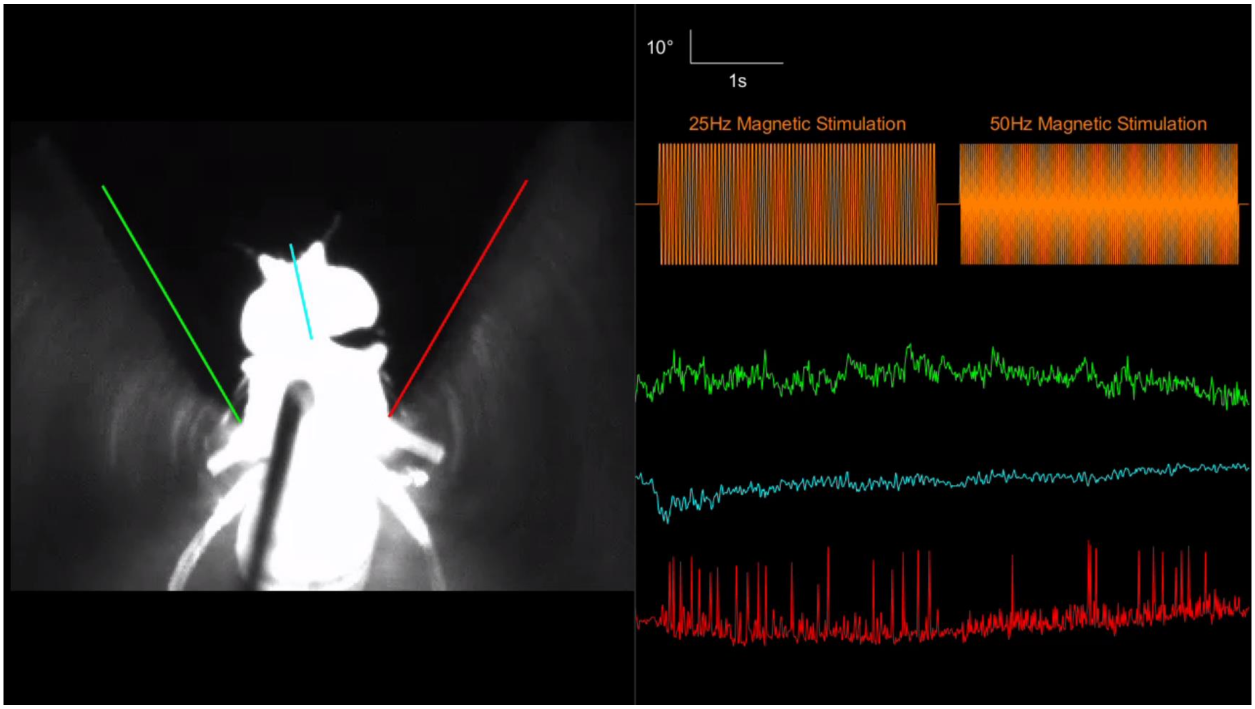


**Supplemental video 2. Kinematic outputs downstream of haltere sensory input**
High speed video (100FPS) showing (from bottom to top) kinematic time series for ipsilateral wing downstroke amplitude (red), head yaw (cyan), contralateral wing downstroke amplitude (green) and interleaved control pulses (orange, video overlay boxes) for magnetic stimulation trials at 25Hz and 50Hz. Note prominent hitches in ipsilateral wing trace (bottom).

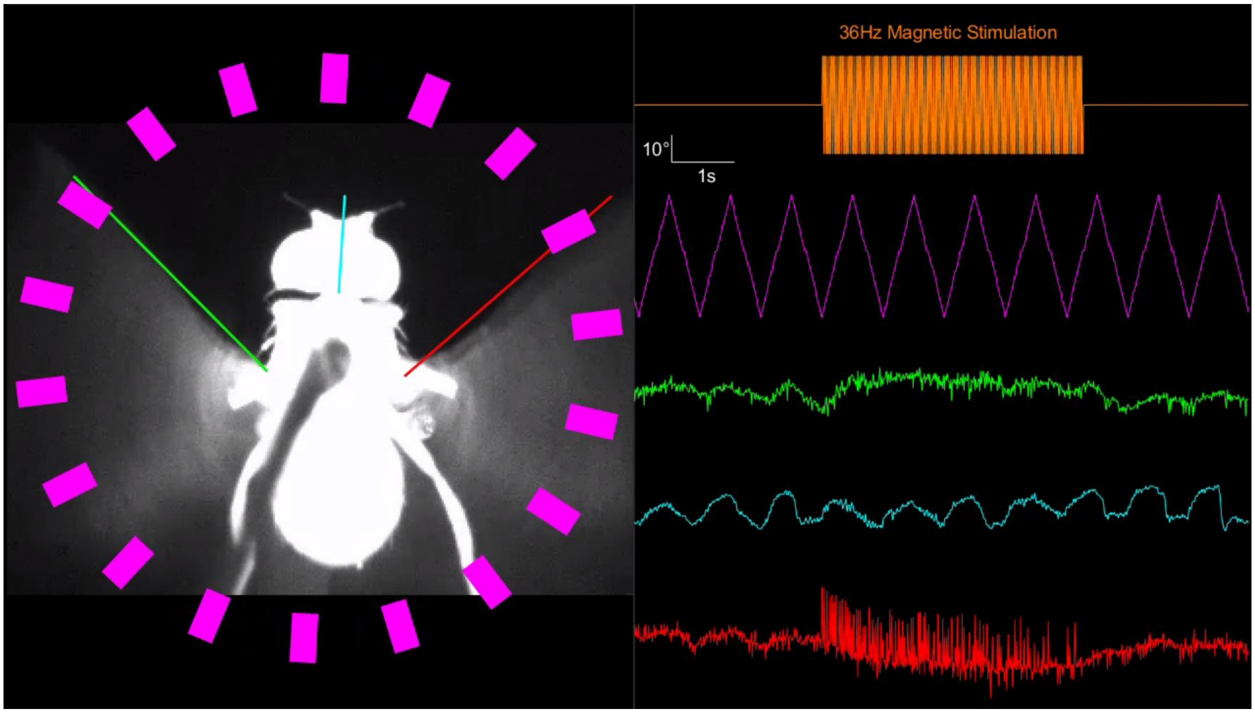


**Supplemental video 3. Concurrent haltere and visual stimulation**
High speed video as in Video S2, showing 36Hz magnetic stimulation of the haltere with concurrent visual motion in the yaw aspect. Reticular magenta overlay and triangle-wave time series (second from top) show angular position of moving stripe visual stimulus. Note cessation of amplitude modulation in contralateral wing (green) during magnetic stimulation epoch.
